# On the neuronal dynamics of aesthetic experience: Evidence from electroencephalographic oscillatory dynamics

**DOI:** 10.1101/2021.06.25.449758

**Authors:** Wim Strijbosch, Edward A. Vessel, Dominik Welke, Ondrej Mitas, John Gelissen, Marcel Bastiaansen

**Author notes:** Academy for Leisure & Events, Breda University of Applied Sciences, Mgr. Hopmansstraat 2, 4817JS Breda, Netherlands.

## Abstract

Aesthetic experiences have an influence on many aspects of life. Interest in the neural basis of aesthetic experiences has grown rapidly in the past decade, and fMRI studies have identified several brain systems supporting aesthetic experiences. Work on the rapid neuronal dynamics of aesthetic experience, however, is relatively scarce. The present study adds to this field by investigating the experience of being aesthetically moved by means of ERP and time-frequency analysis. Participants’ EEG was recorded while they viewed a diverse set of artworks and evaluated the extent to which these artworks moved them. Results show that being aesthetically moved is associated with a sustained increase in gamma activity over centroparietal regions. Also, alpha power over right frontocentral regions was reduced in high and low moving images, compared to artworks given intermediate ratings. We interpret the gamma effect as an indication for sustained savoring processes for aesthetically moving artworks compared to aesthetically less moving artworks. The alpha effect is interpreted as an indication of increased attention for aesthetically salient images. In contrast to previous works, we observed no significant effects in any of the established ERP components, but we did observe effects at latencies longer than 1 s. We conclude that EEG time-frequency analysis provides useful information on the neuronal dynamics of aesthetic experience.

## Introduction

The concept of aesthetic experience has a long tradition within scientific and humanistic literature, with early accounts going back to philosophers such as Baumgarten (1750), Kant (1790) and Hume (1757). This is not surprising, as aesthetic appeal has an influence on many aspects of life, from making purchasing decisions to the enhancement of subjective well-being (see e.g. Cuypers et al., 2012). An aesthetic experience is generally understood as “a perceptual experience that is evaluative, affectively absorbing and engages comprehension (meaning) processes” (Vessel, 2020, p. 1). Aesthetic experiences often have a conceptual component, such as deciphering an abstract work of art. They are typically associated with feelings of pleasure and/or beauty (see e.g. Brielmann & Pelli, 2018; Brielmann & Pelli, 2019), but may encompass more complex responses as well, such as feelings of the sublime or of being moved (Belfi et al., 2019; Menninghaus et al., 2015).

The field of *neuroaesthetics* is involved with examining the neural and behavioral basis of aesthetic experiences in particular (Chatterjee, 2011; Marin, 2015; Pearce et al., 2016; Vessel, 2020). Studies using functional magnetic resonance imaging (fMRI) have identified a number of brain systems that are involved with aesthetic experiences. This circuitry includes sensory and motor pathways, subcortical reward circuitries, the ventromedial prefrontal cortex, and the default-mode network (DMN) (for a detailed review, see Vessel (2020)). In addition to the spatial structure of the aesthetic experience circuitry, the fast temporal dynamics of neural responses to aesthetic experiences have also been examined using both electroencephalography (EEG) and magnetoencephalography (MEG). With respect to the EEG literature in particular, most studies have examined event-related potentials (ERPs) evoked by aesthetically appealing (versus unappealing) stimuli and have sought to link specific ERP components to various stages of aesthetic processing. These efforts (reviewed below) have resulted in substantial insights into the neuronal dynamics of aesthetic experiences, both in terms of their constituent processes (i.e. perception, attention and emotion) and the order in which these processes occur.

Yet beyond these valuable insights from ERP work, other properties of the EEG signal may provide crucial insights for understanding aesthetic experiences as well. This is particularly likely given two key properties of aesthetic experiences: they unfold over a relatively long timescale relative to other cognitive processes (with peak pleasure occurring as late as 3-5 seconds after onset (Belfi et al., 2019; Brielmann, Vale, & Pelli, 2017)) and can be highly idiosyncratic, with different participants responding in very different ways to the same stimulus (Vessel, Maurer, Denker, & Starr, 2018). It is therefore likely that some constituent processes are spread out over longer timescales and not timelocked to stimulus onset. We argue that oscillatory neuronal activity forms a promising approach to better understand the neural basis of aesthetic experiences, as oscillatory dynamics have been effectively related to the coupling and uncoupling of functional networks in the brain (Bastiaansen, Mazaheri, & Jensen, 2012), and can be examined over longer timescales. To date, work on the oscillatory activity related to aesthetic experience remains minimal. In the present study, we thus aim to extensively explore the neuronal dynamics that underly appealing aesthetic experiences.

### EEG and visual aesthetic experience: ERP research

The EEG work on neuroaesthetics has mostly covered visual and auditory stimulus domains. In the current paper, we focus on the work on visual aesthetic experiences. Most EEG studies on visual aesthetic experiences have studied the concept with an event-related approach. Under this approach, various well-established ERP components have been examined in relation to aesthetic experiences.

One ERP component that has been consistently found to be larger for stimuli that are evaluated as aesthetically pleasing is the posterior P2 (Noguchi & Murota, 2013; Righi, Gronchi, Pierguidi, Messina, & Viggiano, 2017). Although its functional properties remain largely unknown (Luck, 2014), the posterior P2 has been previously associated with higher-order perceptual processing modulated by attention (Liu, Meng, Wu, & Huang, 2012; Luck & Hillyard, 1994; Omoto et al., 2010). In this light, an increased posterior P2 for aesthetically pleasing stimuli may suggest that aesthetic appeal prompts a more effective allocation of selective attentional resources in the processing of visual information (Righi et al., 2017). In turn, it could also be that attention drives aesthetic appeal top-down, or that a third factor is driving both.

The N2 and P3 components have been found to be consistently larger for aesthetically appealing images as well (Bölte, Hösker, Hirschfeld, & Thielsch, 2017; De Tommaso et al., 2008; Ma, Hu, & Wang, 2015; Oliver-Rodríguez, Guan, & Johnston, 1999; Righi et al., 2017). Both components have often been found to correlate with one another and have been associated with recognizing target features in stimuli following an experimental task (Luck, 2014). In the neuroaesthetics literature, N2 and P3 effects have only been found in paradigms in which participants were explicitly asked to evaluate stimuli in terms of aesthetic appeal. In paradigms where participants were not asked to do so, there was no difference in the ERPs between aesthetically appealing and unappealing images (Höfel & Jacobsen, 2007a, 2007b), thus suggesting that aesthetic evaluation is an intentional process that does not happen spontaneously.

Nonetheless, the N2 in particular has been used to study the processing order of aesthetic evaluations. The onset of the N2 has been described as “the time by which there must have been enough information available to help the person decide whether or not to respond,” (Schmitt, Munte, & Kutas, 2000, p. 474). Applying this principle, it was found that for artworks, content is processed faster than style, and that for faces gender is processed faster than attractiveness (Augustin, Defranceschi, Fuchs, Carbon, & Hutzler, 2011; Carbon, Faerber, Augustin, Mitterer, & Hutzler, 2018). Righi et al. (2017) found shorter latencies of the N2 for aesthetically appealing stimuli as compared to aesthetically unappealing stimuli, thus suggesting that the information needed to make an aesthetic judgement is earlier at hand when images are aesthetically appealing.

In contrast to the abovementioned effects, the anterior P2 has been found to be consistently larger for aesthetically *unappealing* stimuli (Jiang & Cai, 2013; Ma et al., 2015; Righi et al., 2017; Wang, Huang, Ma, & Li, 2012). Outside the context of aesthetic experience, the anterior P2 has been linked to emotionally negative stimuli as well (Carretié, Mercado, Tapia, & Hinojosa, 2001; Huang & Luo, 2006). However, several other studies have found larger anterior P2 amplitudes for both emotionally positive and negative stimuli compared to neutral (see Hajcak, Weinberg, MacNamara, and Foti (2012) for an overview). Hajcak et al. (2012) thus conclude that the P2 reflects the processing of emotional salience or arousal, rather than the processing of emotionally negative stimuli alone. Larger anterior P2s for emotionally negative stimuli have been connected with the negativity bias: the notion that individuals are especially sensitive to emotionally negative materials, which makes them perceived as more salient (Huang & Luo, 2006).

Results on the P1 and late positive potential (LPP) are less consistent. In some studies, the P1 was larger for aesthetically appealing stimuli (Righi et al., 2017), whereas in other studies it was larger for unappealing stimuli (Bölte et al., 2017) or even indifferent (Noguchi & Murota, 2013). The P1 has been associated with early stages of visual perception based on stimulus characteristics such as color or contrast (Luck, 2014). With the variety of the stimuli that were used in the three aforementioned studies (artworks, websites and everyday objects), it could thus be argued that differences in the P1 were mostly caused by low-level stimulus features, rather than by aesthetic appeal.

In some cases, the LPP was reported to be larger for aesthetically appealing stimuli (Marzi & Viggiano, 2010; Werheid, Schacht, & Sommer, 2007), whereas in other cases it was indifferent between appealing and unappealing stimuli (Jacobsen & Höfel, 2003; Schacht, Werheid, & Sommer, 2008). Schacht et al. (2008), however, found that while there was no difference in the LPP between aesthetically appealing and unappealing stimuli, LPPs for both of the categories were significantly higher than those for images that were evaluated as neutral. The latter is in line with a vast body of work on the LPP (see Hajcak et al. (2012) for review), in which it was found that the LPP is larger for emotionally arousing stimuli, regardless of the emotional valence. Differences in LPPs between aesthetically appealing and unappealing stimuli (Marzi & Viggiano, 2010; Werheid et al., 2007, both in the context of facial attractiveness) may be explained by the notion that attractive faces are generally experienced as more arousing than unattractive faces (North, Todorov, & Osherson, 2010).

### Oscillatory activity and functional networks related to aesthetic experiences

In sum, ERP research has strengthened the notion that aesthetic experiencing encompasses subprocesses of perception, attention and emotion, and that these processes are temporally ordered (e.g. perception of contents before style and perception of gender before facial attractiveness). Other potentially relevant features of the EEG signal, however, have largely been neglected in the research on aesthetic experiences. Oscillatory neuronal activity, for example, is time-but not necessarily phase-locked to an event and especially suited for studying the dynamics in distributed functional networks (Bastiaansen et al., 2012; Varela, Lachaux, Rodriguez, & Martinerie, 2001). Arguably, oscillatory neuronal dynamics may reveal at least some of the temporal dynamics within the network of brain regions that have been identified through fMRI research as important for aesthetic experiences (see above). Notably, as explained in some detail below, oscillatory activity in the theta (4-7 Hz), alpha (around 10 Hz) and gamma (30 Hz and up) frequency ranges can potentially shed light on the temporal dynamics of networks involved in contemplation, attentional allocation and top-down perception, respectively.

Activation of the default-mode network (DMN), a network typically associated with internally directed mental activity as opposed to external focus (Andrews-Hanna, Reidler, Sepulcre, Poulin, & Buckner, 2010; Fox et al., 2005), has been repeatedly linked to the feeling of “being moved” across various aesthetic stimulus domains, ranging from artworks to natural landscapes (Belfi et al., 2019; Vessel, Isik, Belfi, Stahl, & Starr, 2019; Vessel, Starr, & Rubin, 2012, 2013). The DMN generally resides in a suppressed state when perceiving external objects, yet as opposed to other external stimuli, highly moving aesthetic experiences with visual artwork tend to release the DMN from this state of suppression within the first few seconds following stimulus onset (Belfi et al., 2019; Cela-Conde et al., 2013). Activation of the DMN can arguably be related to the contemplative and self-reflective nature of aesthetic processing, although the exact role of the DMN in aesthetic experiences has yet to be established more clearly (Vessel, 2020).

In terms of oscillatory dynamics, activation of the DMN has been related to power decreases in theta-band activity over midfrontal areas. Simultaneous EEG/fMRI recording during resting state shows a negative correlation between midfrontal theta power changes and BOLD signal change in the DMN (Scheeringa et al., 2008). Similar effects were found in EEG/fMRI studies using a working memory task (Meltzer, Negishi, Mayes, & Constable, 2007; Scheeringa et al., 2009). Decreases of theta-band power in midfrontal electrodes thus seem to index DMN activation. Tracking the time course of changes in theta power may therefore be informative for understanding the time course of DMN involvement in moving aesthetic experiences.

A second functional network that might be involved in aesthetic experiencing relates to attention. The ventral attention network has been consistently found to be engaged by aesthetically appealing stimuli across various stimulus domains (Brown, Gao, Tisdelle, Eickhoff, & Liotti, 2011). In turn, attentional networks have frequently been associated with oscillatory activity in the alpha frequency band (Klimesch, 2012). Attentional processes are often characterized by alpha power decreases over task-relevant cortical areas (reflecting increased processing in these areas), together with alpha power increases over task-irrelevant areas (reflecting the suppression of irrelevant information) (Handel, Haarmeier, & Jensen, 2011; Jensen, Bonnefond, & VanRullen, 2012). Specifically, activation of the ventral visual stream (which corresponds to the processing of stimulus contents) has been associated with alpha power decreases over the parietal region in particular (Jokisch & Jensen, 2007). As aesthetically moving stimuli likely engage attention more strongly than non-moving stimuli, we expect that aesthetic appeal corresponds with a decrease in alpha activity over parietal areas.

Third, aesthetic experiences often have a strong conceptual component, such as making sense of an ambiguous work of art or decoding symbolism (Vessel et al., 2019). Arguably, an antecedent of aesthetic evaluation is the active construction of a coherent representation of, say, an artwork from its various elements, such as color, scale and depicted objects (Leder, Belke, Oeberst, & Augustin, 2004). In the empirical aesthetics literature, object recognition has mostly been associated with processes of sense-making (Muth & Carbon, 2013). While it has been hypothesized that familiar and easy processable artworks should be preferred over more difficult alternatives (the so-called fluency hypothesis; see e.g. Reber, Winkielman, and Schwarz (2016)), a growing body of work suggests that in fact ambiguous artworks are preferred as they provide the pleasurable challenge of deciphering patterns and symbols, creating meaning and making sense (Jakesch & Leder, 2009; Muth & Carbon, 2013; Muth, Hesslinger, & Carbon, 2015). Processes of sense-making and constructing object representations have been connected to oscillatory activity in the gamma band (∼40 Hz) at longer latencies following stimulus onset (Bertrand & Tallon-Baudry, 2000; Rodriguez et al., 1999; Tallon-Baudry & Bertrand, 1999). Therefore, we expect that gamma power will increase during the presentation of aesthetically appealing stimuli.

To date, there are very few EEG studies that have examined oscillatory dynamics in a context that comes close to that of an aesthetic experience. Some of these studies have focused on specific channels only and have found suppressed beta activity (Herrera-Arcos et al., 2017) and increased gamma activity (Lopez-Persem et al., 2020) over frontal channels for preferred stimuli compared to non-preferred stimuli. To the best of our knowledge, only one study has examined oscillatory EEG activity across the full scalp (Lindsen, Jones, Shimojo, & Bhattacharya, 2010). Lindsen and colleagues (2010) studied the preference of faces by presenting participants with face pairs (one presented after the other) and subsequently asking them to choose which of the two faces they would prefer approaching. The analysis window was restricted to 200-800 ms following stimulus onset for studying oscillatory activity in five frequency bands: theta (5-8 Hz) alpha (8-12 Hz), beta (12-32 Hz), lower gamma (32-40 Hz) and higher gamma (40-60 Hz). Across the board, there was no difference in oscillatory activity between preferred and non-preferred faces. However, when presentation order was taken into account, preferred faces that were presented second showed increased theta band activity over the left frontal and fronto-central parts of the scalp around 500 ms following stimulus onset. For preferred faces that were presented first, increased gamma activity was observed over central and left parieto-occipital parts of the scalp around 650 ms post-stimulus onset. The gamma effect was interpreted as reflecting an interaction between a relative preference for the first face and a retrieval of its attributes from memory for comparison with the second face. The theta effect was interpreted as a reflecting positive appraisal processes of the chosen face.

Using MEG instead of EEG, Munar and colleagues (2012) studied oscillatory neuronal dynamics in the context of artworks and landscapes. Participants were presented with pictures of landscapes and artworks from various style periods for 3000 ms. For each stimulus, participants were asked to indicate whether they found the stimulus beautiful or not by raising their index finger during stimulus presentation for either one of the two response categories. Oscillatory activity was studied in the time window of 0-1000 ms following stimulus onset in the theta (4-8 Hz), alpha (8-12 Hz), beta (12-30 Hz) and gamma (30-50 Hz) bandwidths. Results showed that from 400 ms onward, there was an increase in activity in all four frequency bands for stimuli that were evaluated as beautiful versus not beautiful. The authors interpreted 1) the theta effect to reflect the coordination of several band networks, 2) the alpha effect to reflect top-down processes (such as expectations or hypothesizing about artworks), 3) the beta effect to reflect supramodal binding and 4) the gamma effect to reflect perceptual feature-binding. Overall, the authors suggested that power increases across the full frequency spectrum might reflect the engagement of an “aesthetic global neuronal workspace” (Munar et al., 2012, p. 9), in which the processing of beautiful stimuli leads to a greater synchronization of neural activity than not beautiful stimuli.

Arguably, the findings are not conclusive: the observations of Lindsen and colleagues (2010) could not be linked to the suggested theta and gamma effects related to DMN activation and top-down object representation, as induced theta and gamma activity at longer latencies could not be observed due to the analysis window of 200-800 ms. However, the findings of Munar et al. (2012) and Lopez-Persem et al. (2020) seem to be in line with the gamma hypothesis as presented in the current study. In addition, Munar et al. (2012) observed a theta effect commencing around 400 ms post-stimulus, although they did not attribute the effect to DMN activity. Contrary to our predictions of a decrease in alpha activity for aesthetically appealing stimuli, Munar et al. (2012) observed an increase. Munar et al. (2012) argue that this effect may reflect the inhibition of top-down driven attention processes. This suggests that aesthetic experiences mostly encompass bottom-up features of attention from 400 ms post-stimulus onset onward. Nonetheless, in Munar et al.’s (2012) study too, the analysis window of 0-1000 ms does not allow for examining how these effects evolve over longer latencies. At the very least, both accounts demonstrate that aesthetically appealing experiences are supported by changes in oscillatory activity across various frequency bands (Lindsen et al., 2010; Munar et al., 2012). Further studying oscillatory dynamics in relation to aesthetic experience thus seems a worthwhile endeavor. In particular, examining these dynamics at latencies beyond 1000 ms in particular seems to be a largely uncharted field.

### Present study

In the present study, we address the paucity in the EEG literature regarding oscillatory activity related to aesthetic experiences. To do so, we presented participants with various stimuli that are known to evoke both aesthetically moving and non-moving experiences (Vessel et al., 2019), and prompted them to report on how aesthetically moving they found them. Following recent developments in empirical aesthetics (Menninghaus et al., 2015), we operationalize aesthetic appeal as “being moved” instead of beauty or attractiveness, as experiences can be aesthetically appealing for reasons other than conventionally defined beauty or attractiveness as well (Vessel, 2020). “Being moved” can serve as an effective summary measure of diverse aesthetic experiences (Vessel et al., 2012), and may thus capture a more encompassing spectrum of the aspects that are associated with them.

EEG-signals were analyzed both in terms of well-established ERP components, and also by examining oscillatory dynamics in a wide frequency range from 1 to 100 Hz. In our ERP analyses, we focused on studying the anterior and posterior P2 and LPP components, as these have previously been linked to the substages of aesthetic experiencing. Based on previous literature, we hypothesized that the anterior and posterior P2 and the LPP components would be larger following aesthetically moving images. Although the N2 and P3 components have also been studied in the context of aesthetic experience, these components have mostly been linked to the experimental paradigm rather than to aesthetic appeal (Höfel & Jacobsen, 2007a, 2007b). Likewise, the P1 has mostly been linked to processing visual stimulus characteristics rather than to aesthetic appeal (Luck, 2014). Although this hampered the formulation of clear hypotheses for these components, we still include them in our analysis in an exploratory manner, as they have not been studied in terms of being aesthetically moved by artworks, but only in terms of aesthetic beauty.

In terms of oscillatory activity, we expect increases in theta and gamma power along with decreases in alpha power for highly moving artworks, but not (or less so) for less moving artworks. Yet given the paucity of existing empirical data, we also explored oscillatory dynamics more broadly, in a wide range of frequencies from 1 to 100 Hz.

## Methods

### Participants

41 first-year students from Tilburg University (17 male, 24 female, age range 18-25) participated in the study and received study credits for their participation. All participating students were right-handed, had normal or corrected to normal vision and no history of neurological disorders. Participants gave their written informed consent in line with the Declaration of Helsinki. All experimental procedures were approved by the Ethics Review Board of the Tilburg School of Social and Behavioral Sciences of Tilburg University (EC-2016.48). Several participants were excluded from further analysis due to excessive artifacts in the EEG recordings (number depending on the respective analysis pipeline; see the section on *Preprocessing* for details).

### Stimulus materials

Stimulus materials consisted of 148 photographs of visual artworks (paintings, collages, woven silks, excluding sculpture) from the Catalog of Art Museum Images Online database (now defunct), as used in a previous study by Vessel et al. (2019). Stimulus materials reflected a variety of periods, styles and genres, and were representative for European, American and Asian cultures. Although part of museum collections, the artworks were chosen such that only lesser-known artworks were part of the stimulus set. Artist’s signatures were removed to avoid recognizability. Paintings were scaled such that they covered 65% of the screen height while maintaining the differences in aspect ratio across the original artworks.

### Procedure

After having read the written instructions and having given their informed consent, participants were familiarized with the lab and mounted with the EEG equipment. After the preparations, they were seated in a dimly lit and sound-attenuating room in front of a computer screen. Participants were asked to stay relaxed and to refrain from excessive head, body and eye movements so as to keep resulting electrical artifacts to a minimum. Participants were instructed to keep their head still while focusing on and evaluating the artworks based on how aesthetically moving they found each of the images. Participants did not need to fix their gaze during stimulus presentation. The images were presented in 10 blocks of 14-15 trials. At the end of each block, participants could take a voluntary mini break. The mini break ended when a participant pressed a button on the keyboard in front of them. Three longer breaks were mandatory (after block 1, 4 and 8), in order to avoid effects of participant fatigue. After all blocks had been presented, the experimenter entered the room and removed the EEG equipment from the participant.

### Design

Each trial began with a fixation cross (black cross on a grey background, relative luminance = 21,59%) presented in the center of the screen for 1000 ms. The artwork image then appeared for 6000 ms. Stimulus presentation time is rather long as compared to standard paradigms for EEG experiments and was selected to allow participants to more fully absorb the image before coming to an evaluation. After the image disappeared, a visual slider bar appeared on the screen and participants were asked to indicate to what extent they felt aesthetically moved on a continuous interval (marked with the anchors “Not at all moving” and “Highly moving” at the left and right ends, respectively). Participants had to position the slider according to their judgement by clicking and dragging it using the mouse and then click the “Next” button in order to confirm their evaluation. A time-out was set at 5000 ms, after which the trial was discarded from further analysis. Clicking the “Next” button initiated a 3000 ms grey screen that served as the intertrial interval.

### EEG recordings

The EEG signals were amplified in a frequency range between DC and 102 Hz, and digitized at a sampling rate of 512 Hz. EEG signals were recorded from 64 locations on the scalp through active Ag-AgCl electrodes (BioSemi, Netherlands), following the extended 10-20 system (Jasper, 1958). Two additional electrodes were placed at the mastoids for offline rereferencing. Another two electrodes served as an electrical recording reference (CMS active electrode) and ground (DRL passive electrode). EOG signals were measured from two bipolar derivations, above and below the left eye for vertical EOG, and from the outer canthi of both eyes for horizontal EOG. Recording parameters were similar to those used for the EEG electrodes as mentioned above.

### Preprocessing of behavioral data

The behavioral data (stimulus evaluations) were coded in a range from −3 for the lower boundary (“not at all moving”) to 3 for the higher boundary (“highly moving”) with in-between intervals of 0.01. These 148 stimulus evaluations were used to group the trials into three response categories (different for each participant): images that were lowly moving (the 37 trials forming the lowest quartile), images that were moderately moving (the 37 trials around the median) and images that were highly moving (the 37 trials constituting the highest quartile). For clarity, we henceforth refer to these three categories as LO, MOD and HI, respectively. We selected the lowest and highest quartiles to serve as LO and HI to ensure the most extreme difference per participant in terms of ratings, as we expect that extremer differences in evaluation ratings between response categories will lead to clearer differences in the ERPs and TFRs.

### EEG data analysis

EEG data were analyzed using Brain Vision Analyzer (Brain Products GmbH, Germany) and the open-source MATLAB-based toolbox FieldTrip (Oostenveld, Fries, Maris, & Schoffelen, 2011). The recorded signals were rereferenced offline to an average of the left and right mastoid channels. A zero phase-shift Butterworth bandpass filter of the 8^th^ order (0.01-100 Hz) was then applied to the data. When individual channels showed excessive artifacts within an otherwise relatively artifact-less recording, these channels were reconstructed using a spherical spline-based topographic interpolation. Data were then subjected to ocular correction procedure based on an independent component analysis (ICA), in order to clean the EEG signals from noise originating from eye movements. From this point onward, preprocessing was split up into two different procedures: one for the ERP analysis and one for the time-frequency analysis, respectively.

For the ERP analysis, data were segmented into trials of 200 ms prestimulus to 6000 ms poststimulus and were baseline-corrected, using the average of the −200-0 ms window. All segments were then visually inspected for eye movement, muscle activity or other artifacts, following a semi-automatic artifact detection procedure. Eight participants were discarded from further ERP analysis because their data contained too many artifacts. Of the remaining participants (*n* = 35, 15 male, 20 female, age range 18-25), only segments containing excessive artifacts were discarded from further analysis (16.2% of all the segments on average). Recordings were then averaged time-locked to stimulus presentation onset (separate averaging for the three different response categories, i.e. LO, MOD and HI). Whenever a segment did not fall into one of these categories, it was omitted from further analysis. The number of remaining segments did not differ between the three response categories (repeated-measures ANOVA between response categories: *F*_2, 60_ = 2.080; *p* = 0.134). This resulted in ERPs at 64 electrode positions for each response category and for each participant. Participant averages were used as input for the statistical analyses (see below). The data were averaged across participants to serve as the input for the graphical representation in Figures 1 and 2.

**Figure 1.**
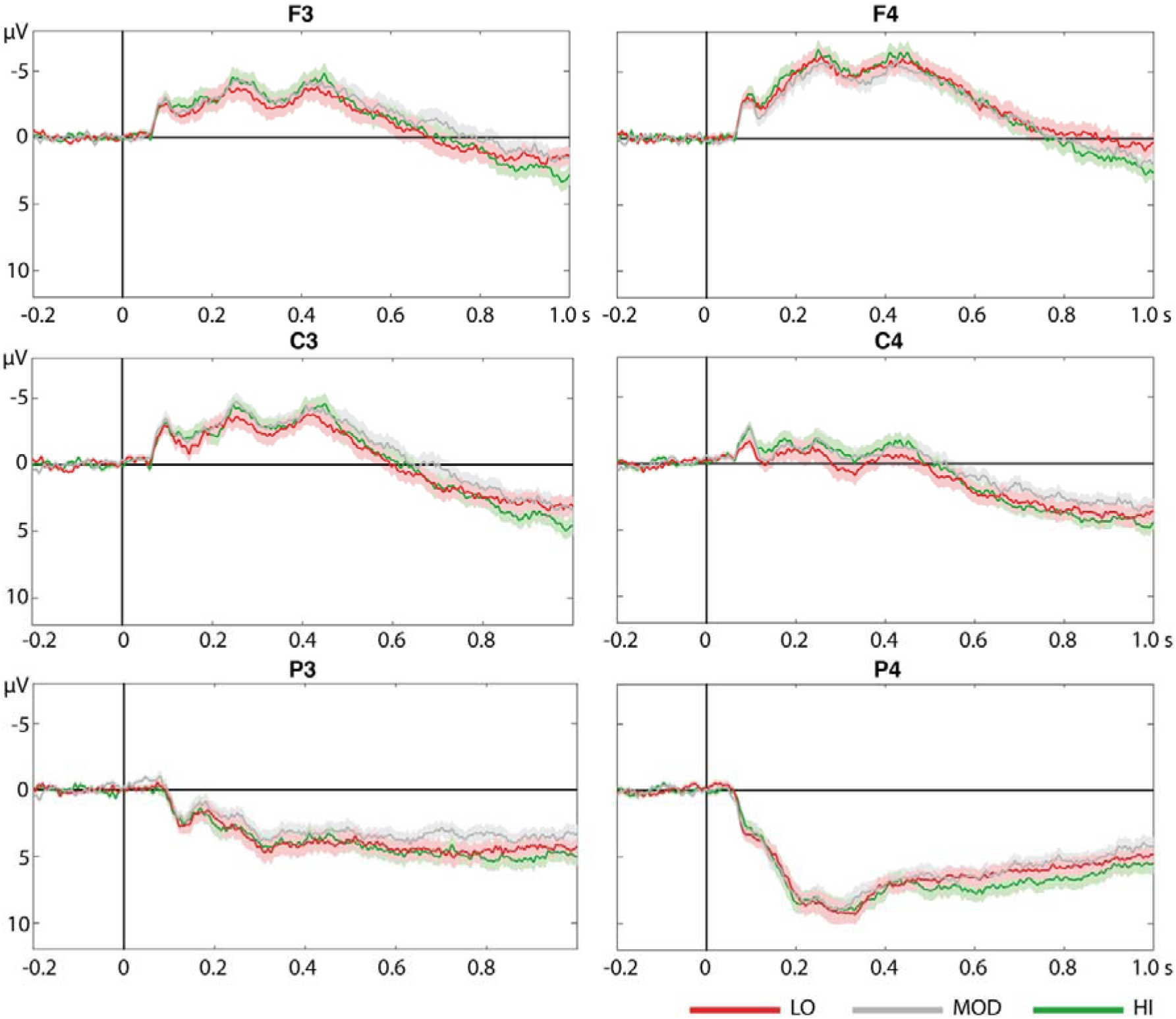
Grand average ERP waveforms (*n* = 35) up to 1000 ms after stimulus onset, evoked by the stimuli in the three different response categories at representative electrodes. Shaded areas around the waveforms indicate standard errors.

**Figure 2.**
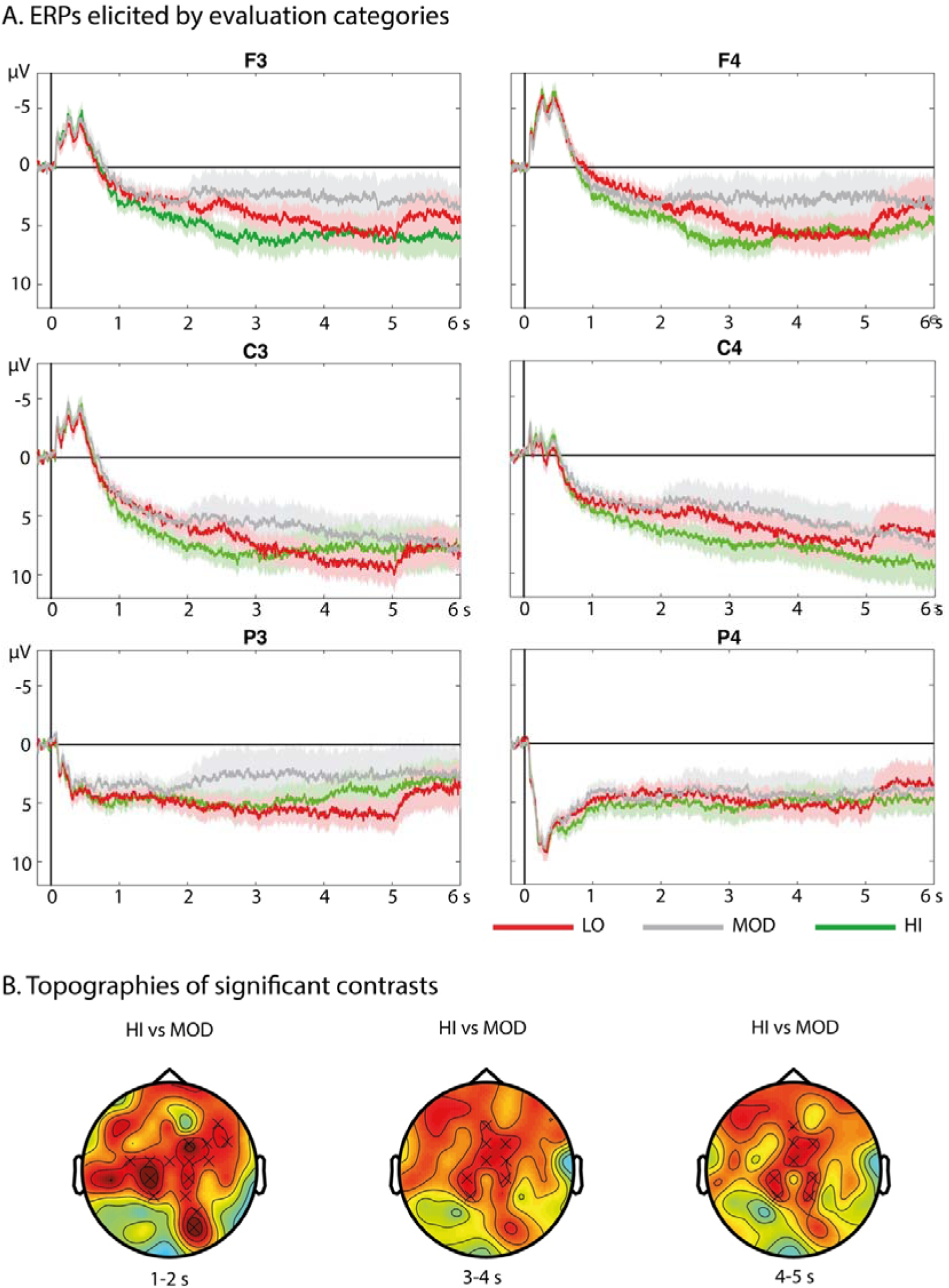
(A) Grand average waveforms (*n* = 31) evoked by the stimuli in the three different response categories at representative electrodes. Shaded areas around the waveforms indicate standard errors. (B) Scalp topographies of the HI versus MOD differences at three different latencies. Electrodes that are part of the significant cluster are marked with an ×.

For the TF analysis, data were first subjected to a second zero phase-shift Butterworth bandpass filter (8^th^ order, 1-100 Hz). Then, data were segmented into trials of 8500 ms around stimulus onset, consisting of a 1500 ms prestimulus interval and a 7000 ms poststimulus interval. The 500 ms prior to stimulus onset served as a baseline. All segments were then visually inspected for artifacts, following a semi-automatic artifact rejection procedure. 10 participants were discarded from further TF analysis because their data contained too many artifacts. Of the remaining participants, (*n* = 31, 14 male, 17 female, age range 18-25), only segments containing excessive artifacts were discarded from further analysis (14.5% of all the segments on average). As the relatively longer trials for TF analysis increase the chance of detecting artifacts, the artifact rejection procedure was slightly more liberal as compared to the artifact rejection procedure for the ERP analysis, leading to a lower percentage of discarded segments. Using the FieldTrip toolbox in MATLAB (Oostenveld et al., 2011), the remaining artifact-free segments were then subjected to two partially overlapping time-frequency transformations: one for lower frequencies (i.e. 2-30 Hz) and one for higher frequencies (i.e. 25-100 Hz). TFRs for lower frequencies were computed using a 400 ms Hanning window, applied in frequency steps of 1 Hz and time steps of 10 ms. TFRs for higher frequencies were computed using a multitaper approach (Mitra & Pesaran, 1999), using a 400 ms time-smoothing and a ±5 Hz frequency-smoothing window, applied in frequency and time steps of 2.5 Hz and 10 ms, respectively. Per participant, the resulting single-trial TFRs were then averaged across trials for the three different response categories (LO, MOD, HI). Whenever a trial did not fall into one of the three categories, it was omitted from further analysis. The number of remaining TFRs did not differ between the three response categories (repeated-measures ANOVA between response categories: *F*_2, 60_ = 1.444; *p* = 0.244). Power changes in the post-stimulus interval were expressed as the relative change to the baseline interval (−500-0 ms) on percentage scale for each time-frequency bin separately. Participant averages were used as input for the statistical analyses. In addition, grand averages across participants were computed for display purposes (see Figure 3 and 4).

**Figure 3.**
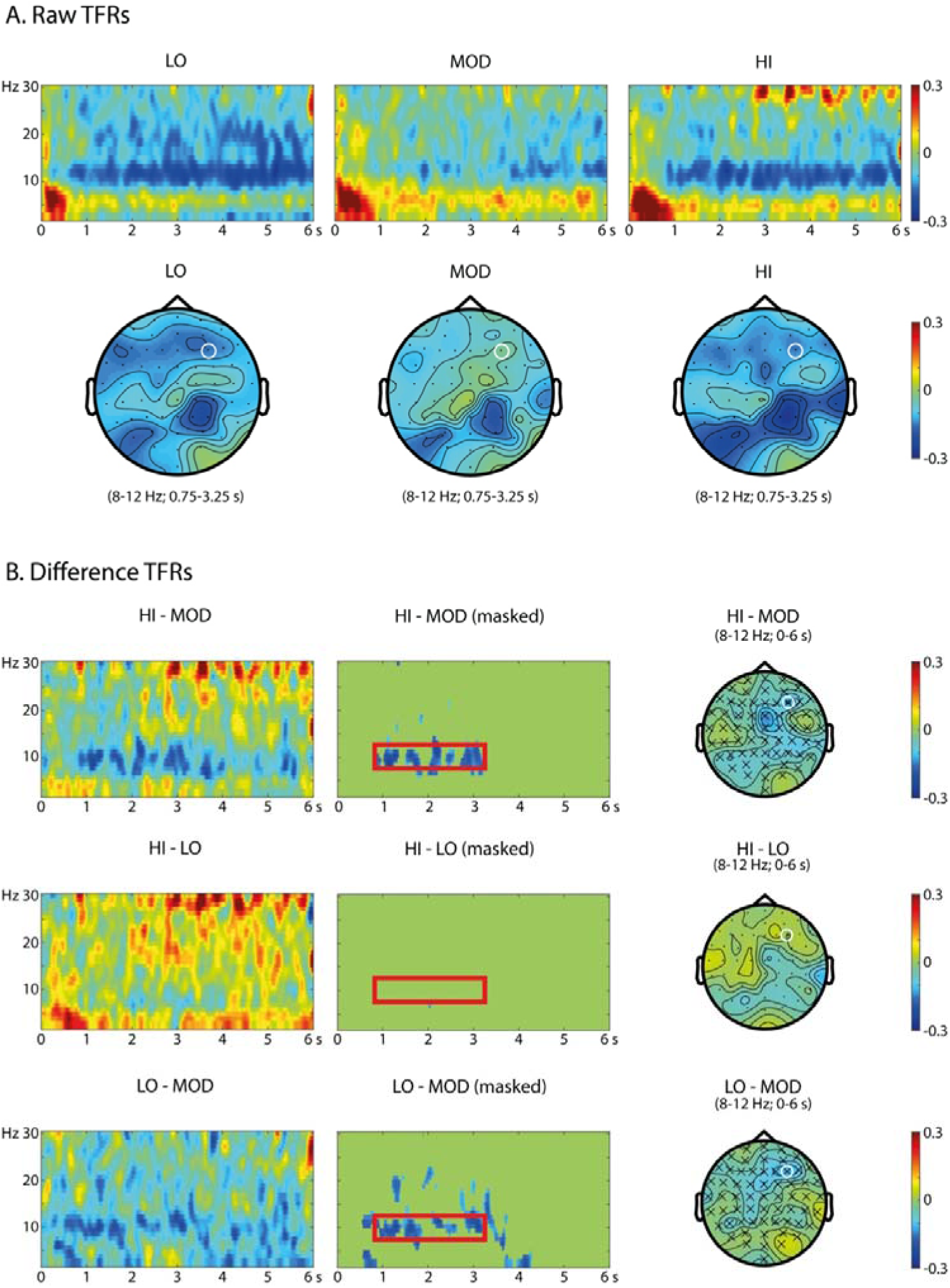
Time frequency representations of single-trial data. (A) Raw TFRs induced by the three response categories in the frequency range of 2-30 Hz. (B) TFR contrasts of HI vs. MOD, LO vs. MOD and LO vs. MOD, with the left panel showing the raw difference, the middle panel showing the masked difference based on the statistical threshold of 5%, and the right panel showing topographic distributions of the observed TFR effects. TFRs are displayed for the F4 electrode (white circle in topo plots). Scales indicate the percentage of signal changes relative to the baseline period (−500-0 ms). Electrodes that are part of the significant cluster are marked with an ×.

**Figure 4.**
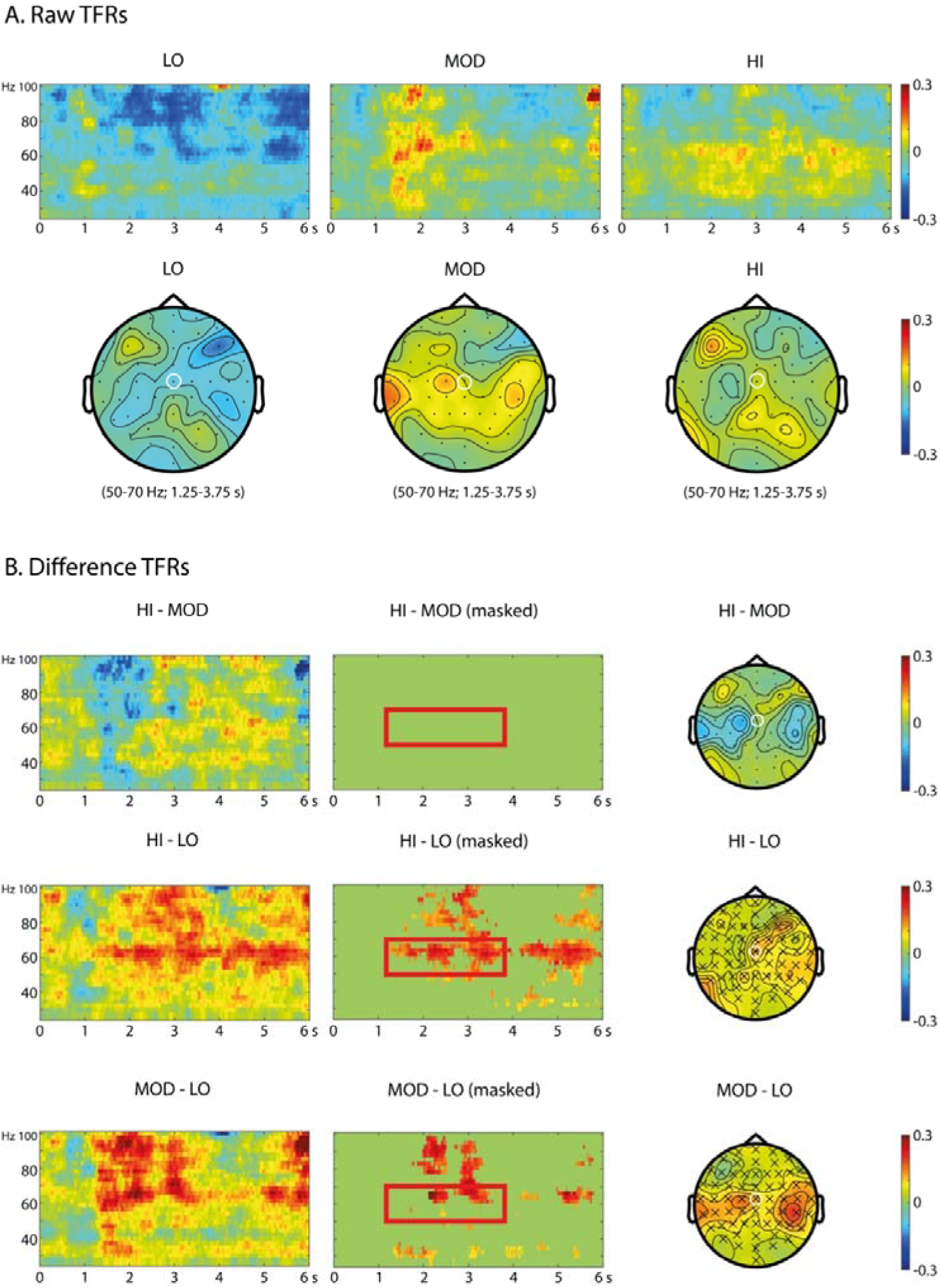
Time frequency representations of single-trial data. (A) Raw TFRs induced by the three response categories in the frequency range of 25-100 Hz. (B) TFR contrasts of HI vs. MOD, HI vs. LO and MOD vs. LO, with the left panel showing the raw difference, the middle panel showing the masked difference based on the statistical threshold of 5%, and the right panel showing topographic distributions of the observed TFR effects. TFRs are displayed for the Cz electrode (white circle in topo plots). Scales indicate the percentage of signal changes relative to the baseline period (−500-0 ms). Electrodes that are part of the significant cluster are marked with an ×.

### Statistical analysis

#### Testing for differences between response categories

To test for significant differences in ERP component amplitudes between LO, MOD and HI, we used a cluster-based random permutation test (Maris & Oostenveld, 2007) for each of the ERP components of interest (anterior and posterior P2 and LPP). In order to explore between-category differences in ERP amplitudes at longer latencies, we performed a test for each subsequent second (i.e. for 1-2 s, for 2-3 s, …, 5-6 s) on the averaged time course of the ERP. To test for significant differences in TFRs, we also used a cluster-based random permutation approach. Tests were performed for the full, non-averaged trial, both in terms of time (0-6 s) and frequencies (2-30 Hz for lower and 25-100 Hz for higher frequencies).

Cluster-based random permutation analysis was used here because it identifies clusters of significant differences between response categories in time, space and frequency, and it elegantly handles the multiple-comparison problem (see Maris and Oostenveld (2007) for a detailed description). For every data point (*electrode × time* for ERP data; *electrode × time × frequency* for TFR data) of two response categories, a dependent-samples *t*-test is performed, yielding uncorrected *p*-values. Clusters of neighboring data points that exceeded the preset α-level of 0.05 are then grouped into clusters. For each cluster thus obtained, a cluster-level test statistic is then defined as the sum of the *t*-statistic in that particular cluster. Then, a null distribution is generated by randomly permuting response categories across participants 1000 times and retaining the largest cluster-level test statistic for each randomization. Finally, the actually observed cluster-level *t*-statistics are compared against this null distribution, and clusters falling in the highest or lowest 2.5^th^ percentile are considered statistically significant.

#### Relating behavioral data to ERP- and TF-based classification accuracy

The distribution of the evaluation ratings was highly different between individual participants – some participants expressed strong opinions about the stimuli, resulting in bimodal distributions of responses near the ends of the slider, while other participants gave less extreme evaluations, generating unimodal distributions near the neutral point on the slider. As such, the relative distances between the three response categories (LO, MOD and HI) were also different across participants. To test whether participants exhibiting more extreme discrimination in self-report between response categories showed more extreme differences in ERPs and TFRs between response categories, we tested whether the difference in self-reported evaluation ratings between the three response categories could predict the classification accuracy of the response categories for the ERP and TFR data. We calculated *d’* measures for each participant between the distributions of evaluation ratings for LO, MOD and HI (i.e. three contrasts in total). *d’*-scores were then correlated with the classification accuracy of the response categories both for the ERP and TF data. Classification was done using an SVM-based classification algorithm with a five-fold cross-validation approach, which was fed with all individual trials per participant (timelocked for the ERP data, and timelocked and TF-transformed for the TF data). Classifications were done for both the full trial (0-6 s), as well as for each separate second in the trial (i.e. for 1-2 s, for 2-3 s, …, 5-6 s). Classification accuracy was defined as the percentage of trials that were correctly classified. To test whether classification accuracies were above chance level, we performed a single-sample *t*-test of the average classification accuracy per category (individual seconds and the full trial) against 50%. For the classification accuracies that were significantly higher than chance level we performed a correlation analysis, reporting Pearson’s correlation coefficient (*r*) and the accompanying *p*-values. As these analyses are secondary to the main analyses on differences in ERPs and TFRs between different response categories of being aesthetically moved, results of these classification analyses are presented in the appendices.

## Results

### Behavioral results

For the trials that remained after artifact correction in the ERP analyses, evaluation ratings ranged from −3 to 0.43 for LO (*M* = −1.73; *SD* = 0.92), from −2.1 to 1.95 for MOD (*M* = 0.20; *SD* = 0.78) and from 0.39 to 3 for HI (*M* = 1.68; *SD* = 0.68). Averages indicate that the evaluation ratings for the MOD category are slightly higher than the neutral point of the scale itself (0). This is also reflected by the behavioral *d’* measures, as the average *d’* for HI vs. MOD (*M* = 3.13; *SD* = 2.04) is slightly smaller than that for LO vs. MOD (*M* = −5.42; *SD* = 2.11) as well.

For the trials that remained after artifact correction in the TF analyses, evaluation ratings ranged from −3 to 0.46 (*M* = −1.72; *SD* = 0.86) for LO, from −1.58 to 1.95 for MOD (*M* = 0.27; *SD* = 0.72) and from 0.37 to 3 for HI (*M* = 1.65; *SD* = 0.67). For these categories too, averages indicate that the evaluation ratings for the MOD category are slightly higher than the neutral point of the scale itself (0), which is again reflected by the behavioral *d’* measures, as the average *d’* for HI vs. MOD (*M* = 3.21; *SD* = 2.08) is slightly smaller than that for LO vs. MOD (*M* = −5.23; *SD* = 2.16).

### ERP results

Figure 1 presents the grand average ERPs for the first 1000 ms after stimulus onset, elicited by the three different response categories for latencies 0-1 s. Figure 2A presents the grand average ERP waveforms for latencies 0-6 s. There were no significant differences between response categories for any of the investigated ERP components. While no hypotheses were formulated for the P3, N2 and P1 components, we rejected the hypotheses on the anterior and posterior P2 and the LPP, for which we expected a difference between highly moving images versus images that were less moving. However, differences were observed for longer latencies. HI generated significantly larger positivity than MOD over anterior-central regions in the time windows of 1-2 s (*p* = 0.024), 3-4 s (*p* = 0.008) and 4-5 s (*p* = 0.018) after stimulus onset. No significant differences were found in the ERP contrasts between LO versus MOD, nor for HI versus LO, although Figure 2 seems to suggest that HI and LO categories consistently elicit larger positivity than MOD through the 1-5 s interval after stimulus onset. Scalp distributions of the observed ERP effects for latencies >1 s are shown in Figure 2B.

### TF results

For the lower frequencies (2-30 Hz), the contrast of HI versus MOD revealed one significant cluster in the frequency range of 8-12 Hz over the right frontocentral region (*p* = 0.022; see Figure 3). This indicates that within the frequency range of 8-12 Hz HI led to a stronger decrease in power than MOD. The contrast of LO versus MOD revealed another contrast in the frequency range of 8-12 Hz over the right frontocentral region (*p* = 0.036; see Figure 3). This indicates that within the frequency range of 8-12 Hz, LO led to a stronger decrease in power than MOD. No differences were found for the contrast of HI versus LO. The observed inverted V-shape provides partial support for our alpha hypothesis, for which we expected a decrease in alpha activity for highly moving images as opposed to images that were less moving.

For the higher frequencies (25-100 Hz), the contrast of HI versus LO revealed one significant cluster in the frequency range of 50-70 Hz over the centroparietal region (*p* = 0.048; see Figure 4). This indicates that within the frequency range of 50-70 Hz, HI led to a stronger increase in power than LO. The contrast of LO versus MOD revealed another significant cluster, also at the frequency range of 50-70 Hz and also over the centroparietal region (*p* = 0.048; see Figure 4). This indicates that within the frequency range of 50-70 Hz, MOD led to a stronger increase in power than LO. We found no significant differences for the contrast of HI versus MOD. These results provide support for our gamma hypothesis, for which we expected an increase in gamma activity for highly moving images as opposed to images that were less moving.

## Discussion

This study aimed to explore the temporal neuronal dynamics of being aesthetically moved by examining the EEG both in terms of oscillations and ERPs. Participants were presented with various artworks and were subsequently asked to indicate how aesthetically moving they found each of the images. Evaluation ratings were grouped into three response categories (LO, MOD and HI) that were then used to study differences in oscillatory neuronal dynamics and ERPs as an effect of being aesthetically highly, moderately or lowly moved. Results indicate that there was an increase in gamma activity for HI and MOD as compared to LO and that there was a decrease in alpha activity for HI and LO as compared to MOD. Contrary to our expectation, we did not observe differences in theta power, nor in any of the ERP components reported before in the context of aesthetic processing.

### Increased gamma activity as an indication of sustained sense-making

We expected gamma activity to increase for aesthetically highly moving compared to less moving artworks. Our data are largely in agreement with this hypothesis, as we observed increased gamma activity (50-70 Hz) over the centroparietal region for HI and MOD as compared to LO. Note that we did not observe significant differences between HI and MOD, which might be due to the fact that the perceived difference in aesthetic appeal between HI and MOD was smaller (*d’* for the ratings = 3.13) than between MOD and LOW (*d’* for the ratings = −5.42). The current findings are in line with the MEG work of Munar et al. (2012), who reported increased gamma activity for artworks that were categorized as beautiful. As increased gamma has been related to processes of visual object representation (Bertrand & Tallon-Baudry, 2000; Tallon-Baudry & Bertrand, 1999), the combined evidence suggests that actively creating a mental representation of an artwork is an important element of the aesthetic experience, which is closely related to evoking feelings of being aesthetically moved.

An additional finding in relation to the increased gamma activity is that it takes some time to develop. Figure 4 shows that for artworks that are aesthetically moving, gamma power increase starts to develop from approximately 1-2 s post-stimulus onset onward and lasts for the full 6 s interval of the analysis window. While the existing studies that addressed oscillatory dynamics related to aesthetic experiences did not analyze their data beyond the first second after stimulus onset (Lindsen et al., 2010; Munar et al., 2012), our findings are in line with previous accounts on object representation (Bertrand & Tallon-Baudry, 2000; Rodriguez et al., 1999; Tallon-Baudry & Bertrand, 1999). The sustained nature of the effect might be characteristic for the aesthetic experience of being moved. In the context of reading poetry, for example, Wassiliwizky, Koelsch, Wagner, Jacobsen, and Menninghaus (2017) found that the feeling of being moved becomes stronger over time, as a poem comes to its close. More broadly, the sustained feeling of being aesthetically moved might be related to the more general concept of savoring: the capacity to attend to, appreciate, and enhance the positive experiences in one’s life (Bryant & Veroff, 2007). Of a variety of experience types, Bryant and Veroff (2007) have suggested that aesthetic experiences are particularly prone to savoring. In particular, we suggest that sense-making might be a prominent feature of savoring that relates to aesthetic experience, which is in line with previous characterizations of aesthetic experience (Jakesch & Leder, 2009; Muth & Carbon, 2013; Muth et al., 2015). Although assessments of aesthetic beauty have been found to be unaffected by stimulus duration (Brielmann et al., 2017), the pleasure associated with beauty has been demonstrated to require thought over time (Brielmann & Pelli, 2017), thus suggesting that savoring or extended sense-making might be an important aspect of aesthetic experiences. The possibility that the presently observed sustained gamma power increases might serve as a neurological proxy for extended sense-making processes corresponding to savoring could form a starting point for future neuroaesthetics research.

### Alpha decreases for aesthetically salient stimuli

Alpha activity was expected to decrease over parietal areas for aesthetically highly moving images compared to less moving images. Contrary to these expectations, we did not observe any significant differences in alpha power changes between HI and LO, although MOD yielded higher alpha power than HI and LO. Also, the modulations in alpha power were found over the right frontocentral region, instead of over parietal regions. The inverted V-shaped alpha power effects as a function of being aesthetically moved are difficult to interpret. Arguably, stronger decreases in alpha power for stimuli that are perceived as either very moving or not moving at all compared to those that are only moderately moving suggest that the inverted V-shaped effect can be interpreted as a salience effect. This interpretation is in line with the fact that alpha is commonly associated with active processing and with attentional processes (Handel et al., 2011; Jensen et al., 2012; Klimesch, 2012). Nonetheless, results are difficult to reconcile with existing data on aesthetic experience and attention (Brown et al., 2011) or attention-related changes in oscillatory alpha power (Munar et al., 2012), both because aesthetic experience has tended to be operationalized as the experience of beauty rather than as being aesthetically moved, and also because non-salient categories have generally not been included (i.e. studies focus on beautiful versus not-beautiful, not on salient versus non-salient categories). In order to be better able to compare results across studies, we therefore suggest that future research in neuroaesthetics should study attention-related changes in alpha power more systematically.

### No effects found for DMN-related theta activity

While the current data show effects for oscillatory neuronal gamma and alpha activity, in contrast with our hypotheses, we did not find any differences in theta power changes between the three response categories. Midfrontal theta power was expected to reflect differential activation of the DMN based on previous fMRI studies that related DMN activity to aesthetic experience (Belfi et al., 2019; Cela-Conde et al., 2013; Vessel et al., 2019; Vessel et al., 2012, 2013) and EEG-fMRI studies that related DMN activity to midfrontal theta power (Meltzer et al., 2007; Scheeringa et al., 2008; Scheeringa et al., 2009). It is unclear at present whether there were indeed no differences in DMN activation in the current study, or whether the absence of midfrontal theta effects suggests that theta power and DMN activity are only linked to each other under the previously established circumstances (i.e. in resting-state activity (Scheeringa et al., 2008) or during working memory tasks (Meltzer et al., 2007; Scheeringa et al., 2009)). Future work would need to address this question more explicitly.

### No differences found for ERP components

Larger anterior and posterior P2 and LPP components were hypothesized for aesthetically highly moving stimuli, as these components have previously been associated with the saliency of an artwork. This hypothesis was not supported by the present findings. Also, we did not find any differences for the P3, N2 and P1 components. We suggest two explanations for the lack of ERP effects in the current study. First, we have used a set of varied, complex stimuli (i.e. artworks from various periods, styles and genres) in combination with a relatively long response window. Previous studies have mostly focused on shorter response windows for less complex sets of stimuli, such as geometric shapes and patterns (De Tommaso et al., 2008; Höfel & Jacobsen, 2007a, 2007b; Jacobsen & Höfel, 2003), faces (Carbon et al., 2018; Lindsen et al., 2010; Marzi & Viggiano, 2010), artworks from only one or two select artists (Augustin et al., 2011) or relatively similar stimuli such as webpages or two-piece suits (Bölte et al., 2017; Jiang & Cai, 2013). Arguably, when using stereotyped, relatively simple stimuli, the evoked mental processes are more homogeneous as well. However, for a more varied and complex set of stimuli, low-level stimulus features may be more diverse, along with the resulting mental processes and their timing. Earlier works on ERPs related to aesthetic experience may therefore have inadvertently focused on mental processing of stimulus-driven differences that were correlated with aesthetic appeal. To support this explanation, further work should aim at systematically comparing the neuronal responses to both complex and less complex sets of stimuli.

A second reason for the lack of an ERP effect could be that thus far, all ERP studies on the aforementioned ERP components in the context of aesthetics have been conducted in experimental paradigms that operationalize aesthetic experience as beautiful/appealing versus non-beautiful/unappealing (Jiang & Cai, 2013; Ma et al., 2015; Marzi & Viggiano, 2010; Noguchi & Murota, 2013; Righi et al., 2017; Wang et al., 2012; Werheid et al., 2007). In line with recent developments in the neuroaesthetics literature (Vessel, 2020), we chose to operationalize aesthetic experience as “being moved” rather than in terms of beauty or attractiveness, as experiences can be aesthetically appealing for reasons other than conventionally defined beauty or attractiveness. The operationalization of aesthetic experience in terms of beauty may lead participants to focus on perceptual features, rather than on other, higher-level features that contribute strongly to being aesthetically moved such as emotion, meaning and self-relevance. In addition, the long response window in our study may allow for more top-down processes to be engaged related to these higher-level processes. Finally, in reviewing the literature on being moved as an aesthetic emotion, Menninghaus et al. (2019) suggest that being moved is an indirect emotion, rather than an emotion that is directly evoked by an aesthetic stimulus. This would be commensurate with a pattern in which effects in the first second post-stimulus onset (i.e. the ERP effects) are absent, but develop subsequently (i.e. the gamma effects and the sustained positivity in the long-latency ERP data). Arguably, aesthetic stimuli trigger general perceptual processes first, which only then contribute to subsequent aesthetic emotions, of which being moved is an exemplary one. However, with the currently available empirical evidence, the above line of reasoning is highly speculative at best.

## Conclusion

This study aimed to explore the rapid neuronal dynamics underlying aesthetic experiences. Using both ERPs and time-frequency analysis, it was found that oscillatory dynamics can be meaningfully linked to aesthetic experiences with a diverse set of visual artworks. In particular, being aesthetically moved was marked by a sustained increase in gamma activity, suggesting that the sustained active construction of mental representations of an artwork has an important role in the experience of being aesthetically moved. In addition, oscillatory alpha power was related to being aesthetically moved through an inverted V-shaped function: alpha power decreased for aesthetically salient stimuli (being highly moved and not being moved at all) but not for non-salient stimuli (being moderately moved). The present findings do not support the hypothesized effects of DMN-related theta power and posterior and anterior P2 and LPP components, nor do they show differences in other previously examined P3, N2 and P1 components. In a broader context, the linking of aesthetics to oscillatory brain dynamics lays the groundwork for identifying the temporally varying component processes of moving aesthetic experiences, and enables further research in more naturalistic settings.

## Acknowledgements

The authors would like to thank Thijs van Laarhoven, Hans Revers and Jeroen Stekelenburg from Tilburg University for their technical help in the laboratory. The authors also would like to thank Andy Brendler and Theresa Hamm for their assistance with the data collection.

## Appendix 1 Predicting ERP classifications from self-report

Classification accuracies for the ERP data for the three different contrasts are presented in Table 1. Results show that of all the contrasts, only the contrast of HI versus LO yielded classification accuracies that were significantly higher than chance level, namely for the full trial (0-6 s) (*p* = 0.009) and for the time intervals of 1-2 s (*p* = 0.008) and 2-3 s (*p* = 0.018). Correlations between these classification accuracies and the respective behavioral *d’* were however not significant (0-6 s: *r* = 0.039, *p* = 0.826; 1-2 s: *r* = 0.042, p = 0.814; and 2-3 s: *r* = 0.188, *p* = 0.287). This indicates that the classification accuracies for HI versus LO for these time intervals was not related to the extremity of the self-reported evaluation ratings between HI and LO.

**Table A1.** Single-sample *t*-tests of ERP-based classification accuracies against chance level

| Classified<br>ERP segment | Contrast |  |  |  |  |  |
| --- | --- | --- | --- | --- | --- | --- |
|  | HI versus MOD |  | LO versus MOD |  | HI versus LO |  |
|  | *M* (*SD*) | *t*<sub>33</sub> | *M* (*SD*) | *t*<sub>33</sub> | *M* (*SD*) | *t*<sub>33</sub> |
| 0 - 6 s | 0,498 (0,082) | -0,115 | 0,511 (0,064) | 0,993 | 0,545 (0,096) | 2,767 |
| 0 - 1 s | 0,518 (0,103) | 0,996 | 0,513 (0,084) | 0,898 | 0,523 (0,073) | 1,866 |
| 1 - 2 s | 0,501 (0,074) | 0,090 | 0,510 (0,077) | 0,730 | 0,539 (0,081) | 2,843 |
| 2 - 3 s | 0,504 (0,103) | 0,221 | 0,524 (0,076) | 1,831 | 0,540 (0,094) | 2,496 |
| 3 - 4 s | 0,485 (0,088) | -0,969 | 0,499 (0,090) | -0,045 | 0,527 (0,099) | 1,582 |
| 4 - 5 s | 0,496 (0,075) | -0,285 | 0,499 (0,075) | -0,073 | 0,519 (0,089) | 1,215 |
| 5 - 6 s | 0,504 (0,072) | 0,317 | 0,499 (0,095) | -0,080 | 0,500 (0,091) | 0,020 |

*Note.* Classification accuracies that are significantly different ( $p < 0.050$ ) from chance level (0.50) are marked in grey.

## Appendix 2 Predicting TF classifications from self-report

Classification accuracies for the TF data for the three different contrasts are presented in Table 3 and 4. Results show that for the frequency range of 2-30 Hz, there are no contrasts that yield classification accuracies that were significantly higher than chance level. For the frequency range of 25-100 Hz, the contrast of HI versus LO did yield classification accuracies that were significantly higher than chance level, namely for the time intervals of 0-1 s (*p* = 0.018) and 5-6 s (*p* = 0.018). Correlations between these classification accuracies and the respective behavioral *d’* were however not significant (0-1 s: *r* = 0.054, *p* = 0.774 and 5-6 s: *r* = −0.273, *p* = 0.137). This indicates that the classification accuracies for HI versus LO for these time intervals could not be related to the extremity of the self-reported evaluation ratings between HI and LO.

**Table A2.** Single-sample *t*-tests of TFR-based classification accuracies against chance level for the frequency range of 2-30 Hz

| Classified<br>TFR segment | Contrast |  |  |  |  |  |
| --- | --- | --- | --- | --- | --- | --- |
|  | HI versus MOD |  | LO versus MOD |  | HI versus LO |  |
| | $M (SD)$ | $t_{30}$ | $M (SD)$ | $t_{30}$ | $M (SD)$ | $t_{30}$ |
| 0 - 6 s | 0,501 (0,100) | 0,074 | 0,490 (0,098) | -0,584 | 0,525 (0,082) | 1,709 |
| 0 - 1 s | 0,495 (0,096) | -0,312 | 0,497 (0,079) | -0,216 | 0,481 (0,085) | -1,249 |
| 1 - 2 s | 0,493 (0,071) | -0,569 | 0,472 (0,086) | -1,829 | 0,521 (0,058) | 2,022 |
| 2 - 3 s | 0,489 (0,077) | -0,814 | 0,509 (0,078) | 0,665 | 0,496 (0,079) | -0,26 |
| 3 - 4 s | 0,470 (0,069) | -2,461 | 0,495 (0,074) | -0,387 | 0,515 (0,069) | 1,167 |
| 4 - 5 s | 0,487 (0,108) | -0,689 | 0,487 (0,072) | -1,022 | 0,496 (0,078) | -0,252 |
| 5 - 6 s | 0,504 (0,084) | 0,244 | 0,500 (0,090) | -0,029 | 0,506 (0,080) | 0,448 |

*Note.* Classification accuracies that are significantly different ( $p < 0.050$ ) from chance level (0.50) are marked in grey.

**Table A3.** Single-sample t-tests of TFR-based classification accuracies against chance level for the frequency range of 25-100 Hz

| Classified<br>TFR segment | Contrast |  |  |  |  |  |
| --- | --- | --- | --- | --- | --- | --- |
|  | HI versus MOD |  | LO versus MOD |  | HI versus LO |  |
|  | *M (SD)* | *t*<sub>30</sub> | *M (SD)* | *t*<sub>30</sub> | *M (SD)* | *t*<sub>30</sub> |
| 0 - 6 s | 0,500 (0,083) | 0,015 | 0,493 (0,081) | -0,498 | 0,529 (0,080) | 2,000 |
| 0 - 1 s | 0,500 (0,088) | -0,028 | 0,492 (0,073) | -0,610 | 0,537 (0,083) | 2,502 |
| 1 - 2 s | 0,513 (0,081) | 0,914 | 0,495 (0,062) | -0,438 | 0,508 (0,083) | 0,538 |
| 2 - 3 s | 0,489 (0,079) | -0,810 | 0,513 (0,089) | 0,822 | 0,523 (0,069) | 1,859 |
| 3 - 4 s | 0,495 (0,083) | -0,329 | 0,519 (0,087) | 1,223 | 0,503 (0,058) | 0,307 |
| 4 - 5 s | 0,516 (0,076) | 1,181 | 0,490 (0,074) | -0,784 | 0,525 (0,075) | 1,874 |
| 5 - 6 s | 0,514 (0,083) | 0,956 | 0,515 (0,083) | 1,004 | 0,539 (0,087) | 2,505 |

*Note.* Classification accuracies that are significantly different ( $p < 0.050$ ) from chance level (0.50) are marked in grey.

